# Remote Activation of a Latent Epitope in an Autoantigen Decoded with Simulated B-Factors

**DOI:** 10.1101/559963

**Authors:** Yuan-Ping Pang, Marta Casal Moura, Gwen E. Thompson, Darlene R. Nelson, Amber M. Hummel, Dieter E. Jenne, Daniel Emerling, Wayne Volkmuth, William H. Robinson, Ulrich Specks

## Abstract

Mutants of a catalytically inactive variant of Proteinase 3 (PR3)—iPR3-Val*^103^* possessing a Ser195Ala mutation relative to wild-type PR3-Val*^103^*—offer insights into how autoantigen PR3 interacts with antineutrophil cytoplasmic antibodies (ANCAs) in granulomatosis with polyangiitis (GPA) and whether such interactions can be interrupted. Here we report that iHm5-Val*^103^*, a triple mutant of iPR3-Val*^103^*, bound a monoclonal antibody (moANCA518) from a GPA patient on an epitope remote from the mutation sites, whereas the corresponding epitope of iPR3-Val*^103^* was latent to moANCA518. Simulated B-factor analysis revealed that the binding of moANCA518 to iHm5-Val*^103^* was due to increased main-chain flexibility of the latent epitope caused by remote mutations, suggesting rigidification of epitopes with therapeutics to alter pathogenic PR3•ANCA interactions as new GPA treatments.

## INTRODUCTION

Proteinase 3 (PR3) is a neutrophil serine protease targeted by antineutrophil cytoplasmic antibodies (ANCAs) in the autoimmune disease granulomatosis with polyangiitis (GPA) (1–5). To investigate how PR3 interacts with the ANCAs during inflammation and whether these interactions can be intervened by therapeutics, we developed a human PR3 mutant (iPR3-Val*^103^*) with a Val*^103^*—the major polymorphic variant at the Val/Ile polymorphic site of the wild-type human PR3 [Val/Ile in GPA patients: 64.6/35.3 (6)]—and a Ser195Ala mutation that alters the charge relay network of Asp102, His57, and Ser195 and thereby disables catalytic functioning in PR3 (7–10). This mutant recognized as many ANCA serum samples from patients with GPA as wild-type PR3 (PR3-Val*^103^*) in both immunofluorescence assay and enzyme-linked immunosorbent assay (ELISA), while the Ser195Ala mutation is close to Epitope 5 of PR3 and remote from Epitopes 1, 3, and 4 as shown in Figure 1 (8, 11). We also developed a number of variants of iPR3-Val*^103^* in the course of our investigation (11).

**Figure 1.**
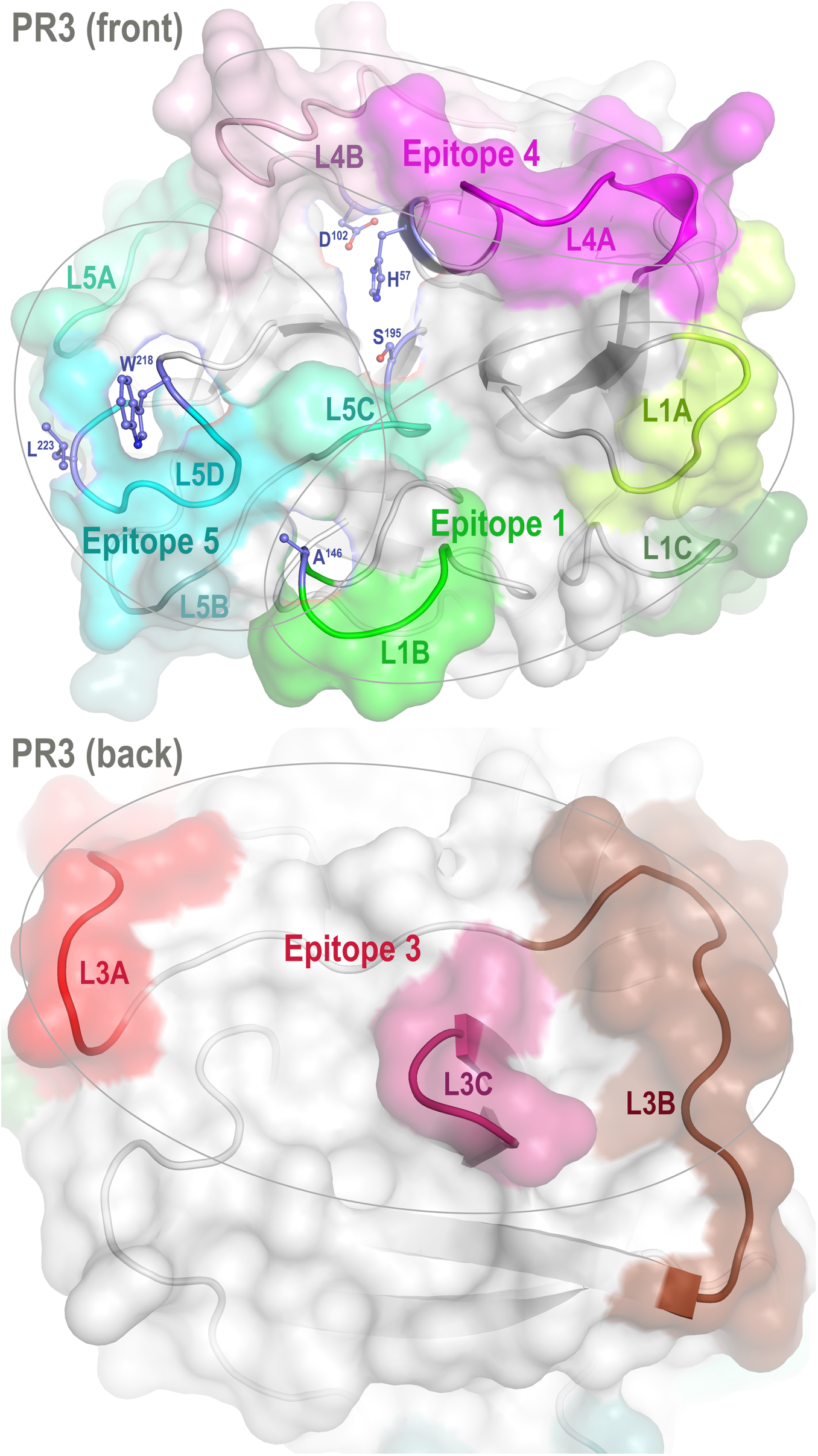
Front and back views of PR3 depicting its four known epitopes, each comprising multiple surface loops with high Cα B-factors derived from simulations. L1A: Loop 1A of residues 36–38C; L1B: Loop 1B of residues 145–151; L1C: Loop 1C of residues 75–79; L3A: Loop 3A of residues 110–117; L3B: Loop 3B of residues 124–133; L3C: Loop 3C of residues 202–204; L4A: Loop 4A of residues 59–63C; L4B: Loop 4B of residues 92–99; L5A: Loop 5A of residues 165–178; L5B: Loop 5B of residues 186–187; L5C: Loop 5C of residues 192–194; and L5D: Loop 5D of residues 219–224; wherein the residue numbering here is identical to that of the PR3 crystal structure (PDB ID: 1FUJ).

One such variant, iHm5-Val*^103^* (formerly referred to as Hm5), has Ala146, Trp218, and Leu223 from human PR3 replaced by Thr146, Arg218, and Gln223 from mouse PR3. Our initial intent of this chimeric triple mutant was to demonstrate reduced binding of ANCAs to Epitope 5 (and possibly Epitope 1 but not Epitopes 3 and 4) of the mutant because Trp218 and Leu223 reside in Epitope 5 and Ala146 is in Epitope 1 as shown in Figure 1 (11). However, as described below, we serendipitously found that a monoclonal ANCA (moANCA518) from a patient with GPA bound to Epitope 3 of iHm5-Val*^103^* but not iPR3-Val*^103^*, although Epitope 3 is distal to the three mutations that reside in Epitopes 1 and 5 (Figure 1). This finding indicates that Epitope 3, a mutation-free epitope of iHm5-Val*^103^*, is latent in iPR3-Val*^103^* but active in iHm5-Val*^103^* for ANCA binding. It also indicates that the latent epitope of PR3 can be activated by remote mutations.

In this context, we raised a mechanistic question: How can a latent antibody binding site in iPR3-Val*^103^* be activated by topologically distal mutations in iHm5-Val*^103^*? The experimental and computational results described below offer insights into this mechanistic question and open a new perspective on a possible cause and novel therapy of GPA.

## MATERIALS AND METHODS

### Materials

Reagents were obtained from Sigma (St. Louis, MO) unless specified otherwise. The human epithelial kidney cell line 293 used for the expression of recombinant PR3 mutants was obtained from ATCC (Rockville, MD).

iPR3-Val*^103^* and iHm5-Val*^103^*: The cDNA constructs coding for iPR3-Val*^103^* and iHm5-Val*^103^* and their expression in HEK293 cells were described in detail elsewhere (11, 12). Both mutants carry a carboxy-terminal cmyc-peptide extension and a poly-His peptide extension for purification using nickel columns from GE Healthcare (Chicago, IL) and for anchoring in ELISAs as previously described and specified below (11–15).

moANCA518: DNA barcode-enabled sequencing of the antibody repertoire was performed on plasmablasts derived from a PR3-targeting ANCA (PR3-ANCA) positive patient as described elsewhere for rheumatoid arthritis and Sjögren syndrome (16–18). Phylograms of the antibody repertoires revealed clonal families of affinity matured antibodies with shared heavy and light chain VJ usage. Twenty-five antibodies were selected for recombinant expression (18) and tested for reactivity with recombinant ANCA antigens [including myeloperoxidase (15), human neutrophil elastase (19–21), iPR3-Val*^103^*, and iHm5-Val*^103^*] using the ELISA. As described in Results, one antibody bound iHm5-Val*^103^* but not iPR3-Val*^103^* and is termed moANCA518, whereas none of the other 24 antibodies bound either of the two PR3 antigens or other ANCA antigens.

Epitope-specific anti-PR3 moAbs: PR3G-2 (22) was a gift from C.G.M. Kallenberg of the University of Groningen. WGM2 (11, 23) was purchased from Hycult Biotech Inc (Wayne, PA). MCPR3-3 was made as previously described (8, 11).

### Enzyme-linked immunosorbent assays

ELISAs used for detection of PR3-ANCA were described in detail elsewhere (12, 13, 15). In brief, either purified PR3 mutants or culture media supernatants from PR3 mutant expressing 293 cell clones diluted in the IRMA buffer (0.05 mM Tris-HCl, 0.1 M NaCl, pH 7.4, and 0.1% bovine serum albumin) were incubated in Pierce® nickel-coated plates from Thermo Fisher Scientific (Waltham, CA) for 1 hour at room temperature; control wells were incubated with the IRMA buffer only. The plates were washed three times with Tris-buffered saline (TBS; 20 mM Tris-HCl, 500 mM NaCl, pH 7.5, and 0.05% Tween 20) in between steps. The ANCA-containing serum samples were diluted 1:20 in TBS with 0.5% bovine serum albumin and incubated in the plates with or without the PR3 mutants for 1 hour at room temperature. The PR3•ANCA complexation was detected after incubation for 1 hour at room temperature with alkaline phosphatase-conjugated goat anti-human IgG (1:10,000 dilution). *P*-Nitrophenyl phosphate was used as substrate at a concentration of 1 mg/mL. The net UV absorbance was obtained by spectrophotometry at 405 nm after 30 minutes of exposure. Similarly, when epitope-specific anti-PR3 moAbs were used to immobilize iHm5-Val*^103^* on Maxisorp® plates from Invitrogen (Carlsbad, CA), complexation of moANCA518 with the antigen was detected after incubation of HRP-conjugated anti-human IgG antibody (1:250 dilution) for 1 hour at room temperature; 3,3’,5,5’-tetramethylbenzidine (Thermo Fisher Scientific®) was used as substrate, and the net UV absorbance was obtained by spectrophotometry at 450 nm after 15 minutes of exposure.

### Western blots

Non-reductive, purified PR3 mutant proteins were loaded (1 µg/lane) onto 12% Tris-HCl gels from BioRad (Hercules, CA). The SDS gel electrophoresis was performed at 180 volts for 35 minutes. The proteins were transferred from gels to nitrocellulose membranes, which were subsequently washed with TBS, blocked for 45 minutes at room temperature with TBS with 0.2% non-fat dry milk. The membranes were then washed twice with TBS with 0.1% Tween 20. Monoclonal antibodies (0.5–1.0 µg/mL) were incubated on the membranes overnight at 4 °C. The membranes were then washed twice with TBS with 0.1% Tween 20 and incubated with goat anti-human or anti-mouse IgG HRP conjugates, diluted to 1:20,000, for 20 minutes at room temperature. The membranes were washed again and developed with the Pierce ECL Western Blotting Substrate kit from Thermo Fisher Scientific (Waltham, MA).

### Statistical analysis

SPSS^®^ Statistics for MacOS, version 25 from IBM (Armonk, NY, USA) was used to calculate the means and standard errors of 3–5 repeat experiments and to compare the means between groups with the two-tailed paired *t*-test.

### Initial conformations of PR3 variants

The initial conformation of PR3-Ile^103^ (residues 16–239; truncated for atomic charge neutrality) was taken from the crystal structure of PR3 (24). The initial conformations of the corresponding PR3-Val^103^ and iPR3-Val^103^ (residues 16–239) were taken from the initial PR3-Ile^103^ conformation with mutations of Ile103Val alone and Ile103Val together with Ser195Ala, respectively. The initial conformation of iHm5-Val^103^ (residues 16–238; truncated for atomic charge neutrality) was taken from the initial PR3-Ile^103^ conformation with mutations of Ala146Thr, Trp218Arg, Leu223Gln, Ile103Val, and Ser195Ala. The crystallographically determined water molecules with residue identifiers of 246–249, 257–259, 261–263, 268, 270, 274–276, 279, 280, 291, 292, 296, 298, 307, 309, and 317 were included in all four conformations. The AMBER residue names of ASP, GLU, ARG, LYS, HID, and CYX were used for all Asp, Glu, Arg, Lys, His, and Cys residues, respectively. All initial conformations were refined via energy minimization using the SANDER module of AMBER 11 (University of California, San Francisco) and forcefield FF12MClm (25) with a dielectric constant of 1.0, a cutoff of 30.0 Å for nonbonded interactions, and 200 cycles of steepest descent minimization followed by 100 cycles of conjugate gradient minimization.

### Molecular dynamics simulations

Each of the four energy-minimized conformations described above was solvated with 5578 (for iHm5-Val^103^) or 5536 (for all other variants) TIP3P (26) water molecules (using “solvatebox PR3 TIP3BOX 8.2”) and then energy-minimized for 100 cycles of steepest descent minimization followed by 900 cycles of conjugate gradient minimization using SANDER of AMBER 11 to remove close van der Waals contacts. The initial solvation box size was 58.268 × 68.409 × 65.657 Å^3^ (for iHm5-Val^103^) or 67.337 × 66.050 × 58.335 Å^3^ (for all other variants). The resulting system was heated from 5 K to 340 K at a rate of 10 K/ps under constant temperature and constant volume, then equilibrated for 10^6^ timesteps under a constant temperature of 340 K and a constant pressure of 1 atm using the isotropic molecule-based scaling. Finally, 20 distinct, independent, unrestricted, unbiased, isobaric–isothermal, 316-ns molecular dynamics (MD) simulations of the equilibrated system with forcefield FF12MClm (25) were performed using PMEMD of AMBER 11 with a periodic boundary condition at 340 K and 1 atm. The 20 unique seed numbers for initial velocities of the 20 simulations were taken from Ref. (27). All simulations used (*i*) a dielectric constant of 1.0, (*ii*) the Berendsen coupling algorithm (28), (*iii*) the particle mesh Ewald method to calculate electrostatic interactions of two atoms at a separation of >8 Å (29), (*iv*) Δ*t* = 1.00 fs of the standard-mass time (25), (*v*) the SHAKE-bond-length constraint applied to all bonds involving hydrogen, (*vi*) a protocol to save the image closest to the middle of the “primary box” to the restart and trajectory files, (*vii*) a formatted restart file, (*viii*) the revised alkali and halide ion parameters (30), (*ix*) a cutoff of 8.0 Å for nonbonded interactions, (*x*) a uniform 10-fold reduction in the atomic masses of the entire simulation system (both solute and solvent), and (*xi*) default values of all other inputs of the PMEMD module. The forcefield parameters of FF12MClm are available in the Supporting Information of Ref. (31). All simulations were performed on a cluster of 100 12-core Apple Mac Pros with Intel Westmere (2.40/2.93 GHz).

### Alpha carbon B-factor calculation

In a two-step procedure using PTRAJ of AmberTools 1.5, the B-factors of alpha carbon (Cα) atoms in PR3 were calculated from all conformations saved at every 10^3^ timesteps during 20 simulations of the protein using the simulation conditions described above except that (*i*) the atomic masses of the entire simulation system (both solute and solvent) were uniformly increased by 100-fold relative to the standard atomic masses, (*ii*) the simulation temperature was lowered to 300 K, and (*iii*) the simulation time was reduced to 500 ps. The first step was to align all saved conformations onto the first saved conformation to obtain an average conformation using the root mean square fit of all Cα atoms. The second step was to perform root mean square fitting of all Cα atoms in all saved conformations onto the corresponding atoms of the average conformation. The Cα B-factors were then calculated using the “atomicfluct” command in PTRAJ. For each protein, the calculated B-factor of any atom in Table S2 was the mean of all B-factors of the atom derived from 20 simulations of the protein. The standard error (SE) of a B-factor was calculated according to Eq. 2 of Ref. (32). The SE of the average Cα B-factor of each PR3 variant was calculated according to the standard method for propagation of errors of precision (33). The 95% confidence interval (95%CI) of the average Cα B-factor was obtained according to the formula mean ± 1.96×SE because the sample size of each PR3 variant exceeded 100.

### Conformational cluster analysis and root mean square deviation calculation

The conformational cluster analyses were performed using CPPTRAJ of AmberTools 16 with the average-linkage algorithm (34), epsilon of 3.0 Å, and root mean square coordinate deviation on all Cα atoms of the proteins (Table S1). Cα root mean square deviations (CαRMSDs) were manually calculated using ProFit V2.6 (http://www.bioinf.org.uk/software/profit/). The first unit of the crystal structure of the PR3 tetramer and the time-averaged conformation (without energy minimization) of the most populated cluster were used for the CαRMSD calculation and Figure 3.

## RESULTS

In characterizing moAbs identified and cloned from B cells in patients with GPA, we found that one of these, moANCA518, bound to iHm5-Val*^103^* but not iPR3-Val*^103^* (Figure 2A) according to the ELISA using iHm5-Val*^103^* and iPR3-Val*^103^* both of which contain a *C*-terminal poly-His tag for anchoring the antigens without perturbing the folded conformations of the antigens and without blocking the epitopes of the antigens (12). Further, the binding of moANCA518 to iHm5-Val*^103^* was dose dependent (Figure 2A) and confirmed by the Western blot under non-reducing conditions (Figure S1) as well as by ELISAs using untagged PR3 variants (data not shown). This serendipitous finding prompted us to investigate how the triple chimeric mutations in iHm5-Val*^103^* changed the conformation of iPR3-Val*^103^* and consequently the antigenicity to moANCA518.

**Figure 2.**
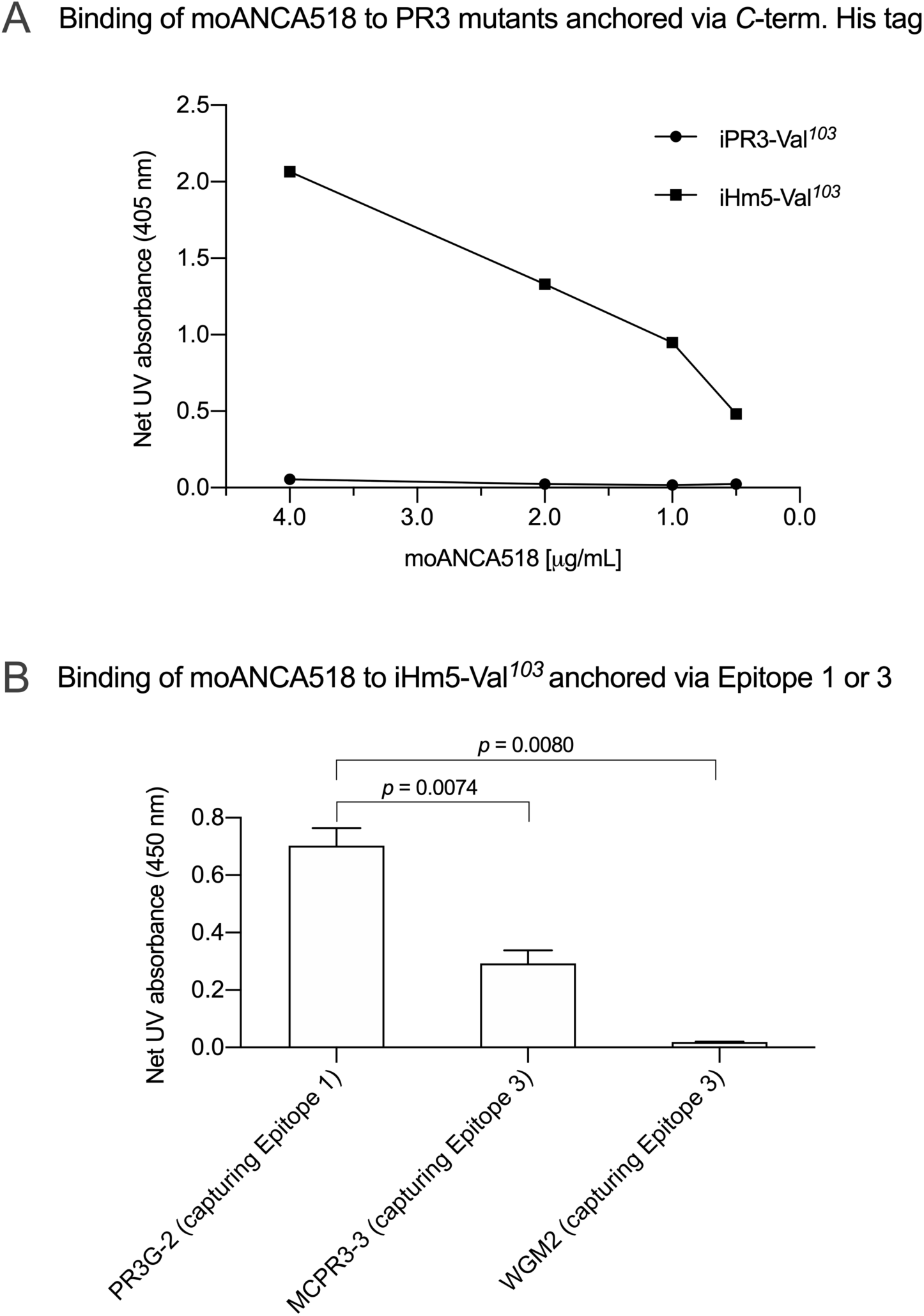
Selective binding of moANCA518 to Epitope 3 of iHm5-Val*^103^*. **A.** Dilution curves show dose-dependent binding of moANCA518 to iHm5-Val*^103^* (solid line) but not iPR3-Val*^103^* (dashed line) in the ELISA using an antigen whose C-terminal poly-His tag is anchored at the plate. The culture media supernatants from PR3 mutant expressing 293 cells were used in the ELISA. **B.** Epitope-specific anti-PR3 moAbs PR3G-2, MCPR3-3, and WGM2 (2, 4, and 4 µg/mL, respectively), which were coated to the plate and used to capture iHm5-Val*^103^* in the ELISA, show Epitope 3 of iHm5-Val*^103^* as a major target site by the primary antibody moANCA518 (1.0 µg/mL). The purified PR3 mutants were used in the ELISA.

Accordingly, we developed computer models of PR3-Val*^103^*, iPR3-Val*^103^*, and iHm5-Val*^103^* to understand how mutations of these variants affect the ANCA-binding capabilities of the four reported epitopes of PR3 (11). These models were derived from MD simulations using our published forcefield and simulation protocol (25), which reportedly folded fast-folding proteins in isobaric–isothermal MD simulations to achieve agreements between simulated and experimental folding times within factors of 0.69–1.75 (35) and are hence suitable for predicting in vivo conformations of PR3 and its variants. The initial conformations of the three variants used in these simulations were derived from the PR3-Ile*^103^* crystal structure (24) because experimentally determined structures of these variants have been unavailable to date. Although small differences in the time-averaged main-chain conformations of two surface loops (Loops 3 and 5) between iHm5-Val*^103^* and PR3-Val*^103^* (or between iHm5-Val*^103^* and iPR3-Val*^103^*) were observed (Figure 3), the overall conformations of the three variants resembled one another according to the Cα root mean square deviations of ≤1.63 Å (Table S1). Given these conformational properties, we could not determine how mutations of these variants affect the ANCA-binding capabilities of the PR3 epitopes, primarily because these surface loops are highly flexible and lack the time dimension (due to time-averaging) that is required for immunological function analysis (36).

**Figure 3.**
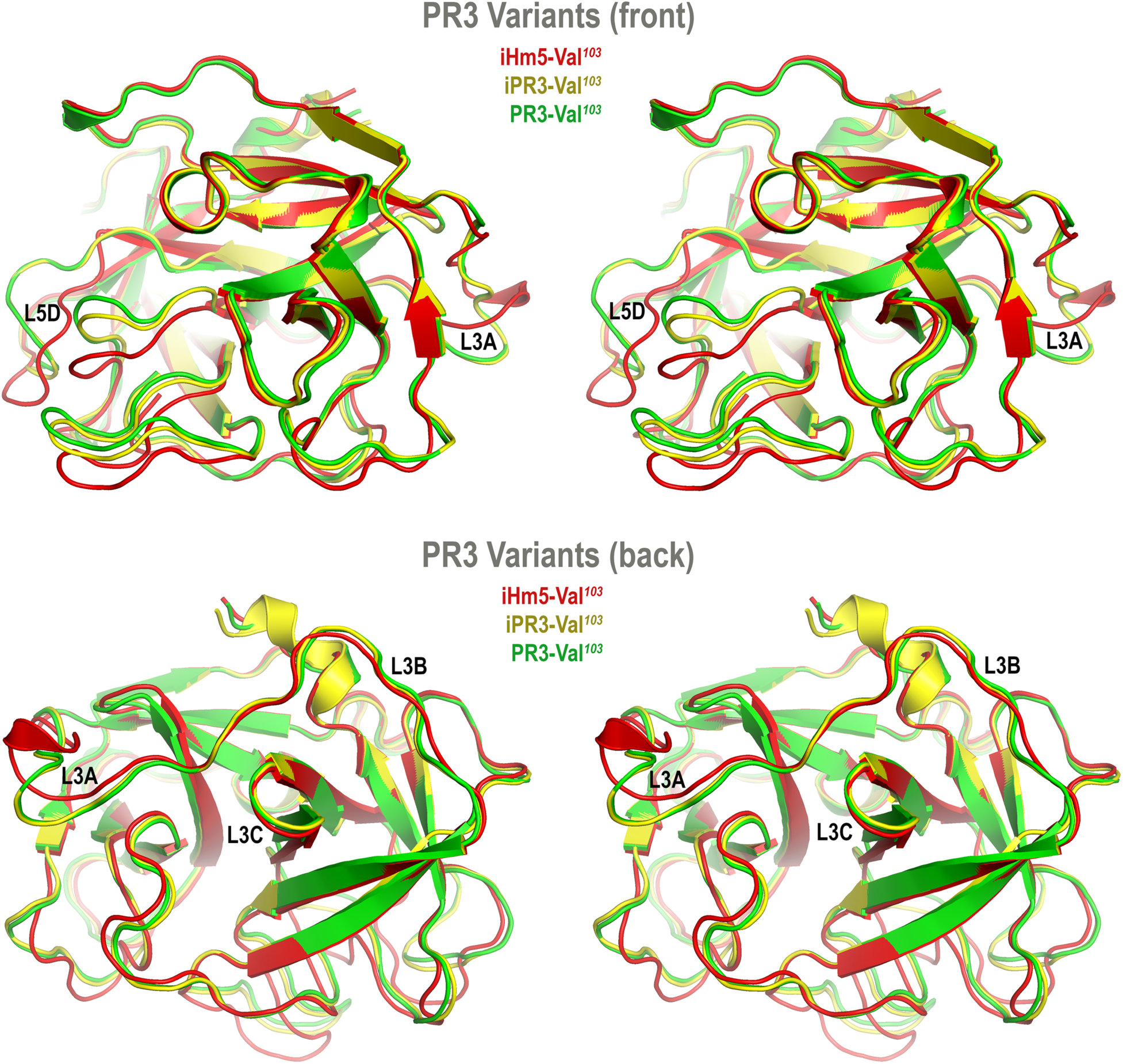
Superimposed time-averaged conformations of three PR3 variants in cross-eye stereo view. The time-averaged conformations were obtained via cluster analyses (without energy minimization) from the most populated cluster in the three sets of molecular dynamics simulations. The L3A in iHm5-Val***^103^*** is visibly structured to a helical component relative to that in two other variants, indicating that L3A in iHm5-Val***^103^*** is less mobile than that in the others. The L3B in iHm5-Val***^103^*** is slightly contracted (due to time-averaging) relative to that in two other variants, indicating that L3B is more mobile than that in the others. See Figure 4 legend for definition of L3A, L3B, L3C, and L5D.

To take the time dimension into account, we turned our attention to the dynamic properties of the PR3 variants. It is well-known that a folded protein is fluid-like with fluctuations in atomic position on the picosecond timescale and that the dynamics of these atomic displacements are dominated by collisions with neighboring atoms involving reorientation of side chains or localized portions of the backbone (37). Two seminal studies have also shown that the crystallographically determined high B-factors of a protein fragment are linked to the antigenicity of the fragment (38, 39). This link indicates that the crystallographically determined B-factor—defined as 8π^2^〈*u*^2^〉 to reflect the displacement *u* of the atom from its mean position, thermal motions, local mobility, or the uncertainty of the atomic mean position (40–48)—can be used to aid the identification and characterization of epitopes.

However, the crystallographically determined B-factor of an atom reflects not only the thermal motion or local mobility of the atom but also conformational and static lattice disorders of the atom, and even the refinement error in determining the mean position of the atom (43, 45, 47, 49). Therefore, using crystallographically determined B-factors to investigate epitopes requires the comparison of B-factors of different crystal structures of the same protein, which are in different space groups and obtained with different refinement procedures at different resolutions, in order to identify the B-factors that reflect the local mobility of the protein (49).

This requirement can be avoided by using simulated B-factors derived from MD simulations on a picosecond timescale because simulated B-factors are devoid of refinement errors and conformational and static lattice disorders. In addition, local motions, such as those of backbone N–H bonds, occur on the order of tens or hundreds of picoseconds (50).

In this context, we calculated the Cα B-factors of PR3-Val*^103^*, iPR3-Val*^103^*, and iHm5-Val*^103^* from MD simulations on a 50-ps timescale using our published forcefield (25) and method (51). The mean Cα B-factors of PR3-Val*^103^*, iPR3-Val*^103^*, and iHm5-Val*^103^* were 6.84 Å^2^ (95%CI: 6.75–6.94 Å^2^), 6.91 Å^2^ (95%CI: 6.82–7.00 Å^2^), and 7.13 Å^2^ (95%CI: 7.03–7.24 Å^2^), respectively. Given these findings, we concluded that any surface loop is highly mobile and hence potentially antigenic if the mean Cα B-factor of the loop was >9.00 Å^2^. This conservative cutoff of 9.00 Å^2^ was based on the mean Cα B-factors of all PR3 variants used in this study (6.84, 6.91, and 7.13 Å^2^). According to this criterion, PR3-Val*^103^* has 10 potentially antigenic surface loops, and iPR3-Val*^103^* and iHm5-Val*^103^* have 11 each (Figure 4). Consistent with the two seminal reports (38, 39), all of these potentially antigenic loops identified a priori by using simulated B-factors fall within all four known epitopes of PR3 (11), demonstrating a clear association between a loop with a high mean simulated Cα B-factor and the experimentally determined antigenicity of the loop.

**Figure 4.**
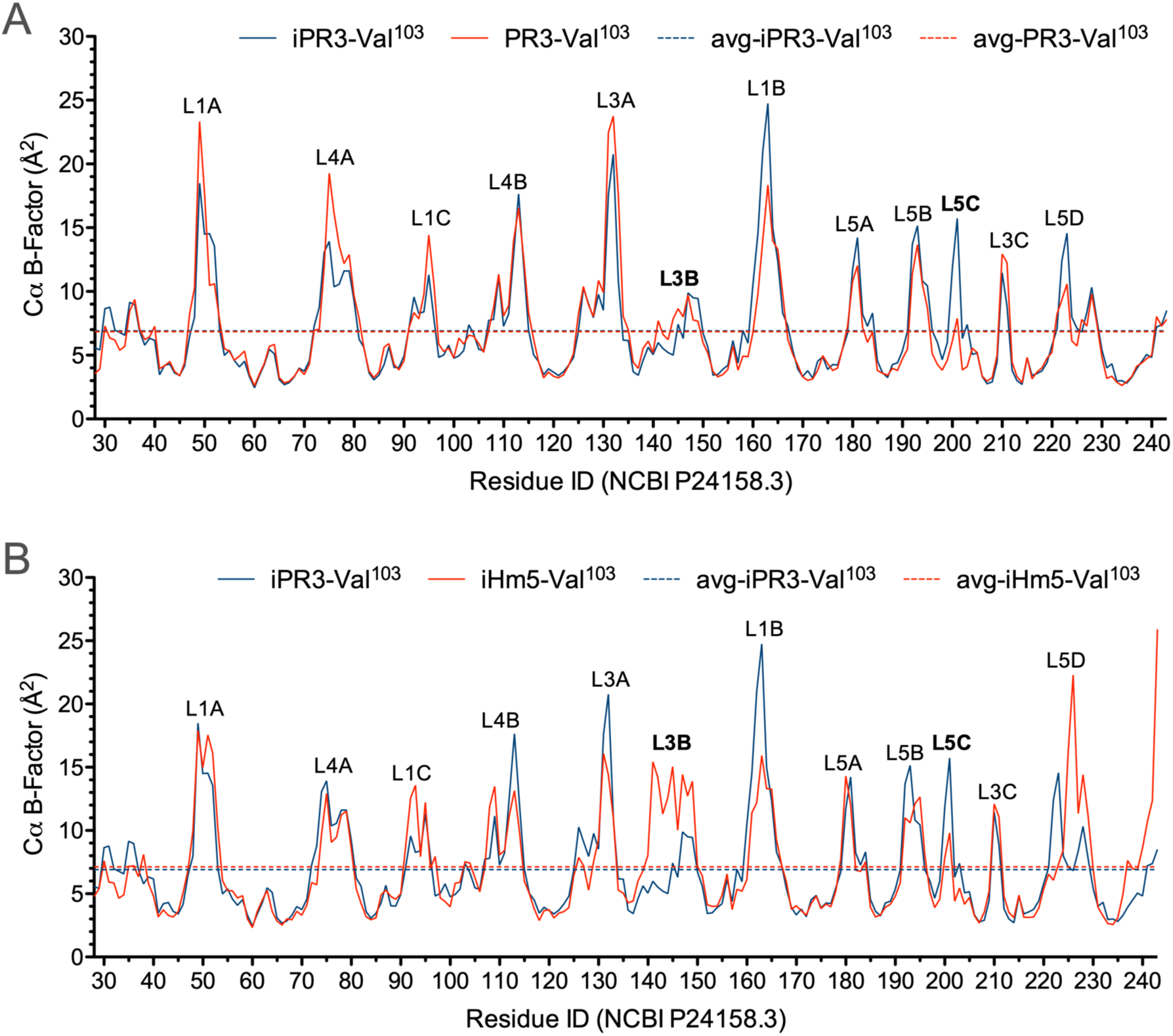
Simulated Cα B-factors of PR3-Val*^103^*, iPR3-Val*^103^*, and iHm5-Val*^103^*. The simulated mean Cα B-factors of PR3-Val***^103^***, iPR3-Val***^103^***, and iHm5-Val***^103^*** are 6.84 Å^2^ (95%CI: 6.75–6.94 Å^2^; labeled as avg-PR3-Val***^103^***), 6.91 Å^2^ (95%CI: 6.82–7.00 Å^2^; labeled as avg-iPR3-Va***^103^***), and 7.13 Å^2^ (95%CI: 7.03–7.24 Å^2^; labeled as avg-iHm5-Val***^103^***), respectively, wherein 95%CI is the abbreviation of 95% confidence interval. The simulated Cα B-factors were plotted using the human PR3 sequence (NCBI P24158.3) numbering because the PR3 crystal structure numbering is discontinuous. Therefore, the following loop residues are defined using the PR3 crystal structure numbering followed by the NCBI P24158.3 numbering in parenthesis. L1A: Loop 1A of residues 36–38C(48–52); L1B: Loop 1B of residues 145–151(161–166); L1C: Loop 1C of residues 75–79(92–96); L3A: Loop 3A of residues 110–117(126–133); L3B: Loop 3B of residues 124–133(140–149); L3C: Loop 3C of residues 202–204(210–212); L4A: Loop 4A of residues 59–63C(73–80); L4B: Loop 4B of residues 92–99(108–115); L5A: Loop 5A of residues 165–178(180–184); L5B: Loop 5B of residues 186–187(192–195); L5C: Loop 5C of residues 192–194(200–202); L5D: Loop 5D of residues 219–224(223–229).

Further, we found that the Ser195Ala mutation caused no significant reduction in the mean Cα B-factor of any of the 10 potentially antigenic surface loops in PR3-Val*^103^* (Figure 4A). This finding implies that the Ser195Ala mutation does not impair the ANCA-binding capability of any of the four epitopes of iPR3-Val*^103^*, and it explains our reported observation that iPR3-Val*^103^* recognizes as many ANCA serum samples as PR3-Val*^103^* does (8).

We also found the mean Cα B-factors of Loop 3B in iPR3-Val*^103^* (possessing Ala146, Trp218, and Leu223) and iHm5-Val*^103^* (possessing Thr146, Arg218, and Gln223) to be 6.9 Å^2^ (95%CI: 6.8–7.0 Å^2^) and 12.8 Å^2^ (95%CI: 12.3–13.2 Å^2^), respectively (Figure 4B). According to the afore-described antigenicity criterion of 9.00 Å^2^, these means suggest that the three chimeric mutations make Loop 3B (a mutation-free loop) more mobile in iHm5-Val*^103^*, despite large separations between Epitope 3 of PR3 and the chimeric mutation sites (∼32 Å, ∼32 Å, and ∼31 Å from the Cα atom of Gln122 in Epitope 3 to the Cα atoms of Ala146, Trp218, and Leu223, respectively, at the chimeric mutation sites). The higher mobility of Loop 3B in iHm5-Val*^103^* relative to that in iPR3-Val*^103^* is also evident from the slight loop contraction (due to time-averaging) of Loop 3B in iHm5-Val*^103^* shown in Figure 3. Therefore, Epitope 3 of iHm5-Val*^103^* could bind ANCAs, whereas the ANCA-binding capability of Epitope 3 of iPR3-Val*^103^* would be rather limited.

We subsequently repeated the afore-described ELISAs in the presence of epitope-specific moAbs that target either Epitope 1 or 3 of PR3. Consistently, we found that PR3G-2 that targets Epitope 1 of PR3 (22) did not affect the binding of moANCA518 to iHm5-Val*^103^*, whereas MCPR3-3 and WGM2, both of which recognize Epitope 3 of PR3 (11), reduced and abolished the moANCA518 binding (*p* < 0.01; Figure 2B), respectively. We also confirmed the binding of moANCA518 primarily to Epitope 3 of iHm5-Val*^103^* using Fabs from epitope-specific moAbs that target Epitope 2 or 5 of PR3 (8, 11, 52) (data not shown).

## DISCUSSION

In view of the data above, we suggest a new mechanism for epitope activation of PR3: Remote mutations can increase the local mobility (i.e., main-chain flexibility) of a latent epitope of PR3, which facilitates the conformational adaptation required for antibody binding and thereby activate the latent epitope. This type of exquisite epitope activation—achieved either in vitro by remote mutations as we demonstrated or in vivo conceivably by remote polymorphisms or by remote protein•ligand binding including allosteric binding with an autoantibody—may be a fundamental feature of GPA. There is evidence that increased mobility of Epitope 3 occurs in vivo as more than 50% of serum samples from patients with GPA preferentially bind iHm5-Val*^103^* (53). It is worth noting that the latent epitope activation described here conceptually differs from the exposure of cryptic epitopes caused by citrullination (*viz.,* post-translational conversion of arginine to citrulline) (54). The latent epitope activation is due to the significant increase of main-chain flexibility of Loop 3B shown in Figure 4B caused by the three remote mutations, whereas the cryptic epitope exposure is reportedly due to conformational changes triggered by multiple citrullinations (54). The latent epitope activation does not involve conformational changes as the remote mutations do not significantly change the main-chain conformation of Loop 3B shown in Figure 3B. It is also worth noting that identifying PR3 mutations in patients with GPA that can increase the Epitope 3 mobility is difficult because other factors such as remote protein•ligand interactions may also increase the latent epitope mobility in vivo, namely, it is challenging to identify the cause of the latent epitope activation in vivo.

Nevertheless, knowing the increased mobility of Epitope 3 of iHm5-Val*^103^* responsible for its binding to moANCA518 alone may have implications for the development of novel, effective treatments of GPA that aim to disrupt the pathogenic autoantibody•autoantigen interactions in GPA by reducing the mobility of epitopes targeted by PR3-targeting ANCAs (PR3-ANCAs). For example, the present finding may explain in principle why a monoclonal antibody strategy (that targets native PR3 and prevents binding of pathogenic PR3-ANCAs to the PR3 that is not in itself pathogenic) is of advantage for disrupting the autoantibody•autoantigen interactions over the molecular decoy strategy (that targets pathogenic autoantibodies). For the latter, large numbers of decoys are required to block a stock of distinct, pathogenic PR3-ANCAs. The DNA recombination and affinity maturation mechanisms, which create diversity and potency in specificity of antibodies, can potentially lead to resistance against the decoys. For the former, only one or a few small-molecule or protein (e.g., monoclonal antibody) binders are required to rigidify B-cell epitopes of PR3 and consequently make the autoantigen inaccessible to a repertoire of distinct, pathogenic PR3-ANCAs, thus obviating mechanisms that could potentially lead to resistance against such binders.

## AUTHOR CONTRIBUTIONS

D.R.N. and U.S. initiated the collaboration project. U.S. and D.E.J. designed the PR3 variants and ANCA-binding experiments. M.C.M., G.E.T., A.M.H., and D.R.N. performed ANCA-binding experiments. Y.-P.P. designed and performed B-factor calculations. D.E., W.V., and W.H.R. discovered moANCA518. Y.-P.P., U.S., and D.E.J. wrote the manuscript. All authors contributed to revisions of the manuscript.

## ACKNOWLEDGEMENT

The authors wish to thank Dr. C.G.M. Kallenberg of the University of Groningen for providing us with epitope-specific anti-PR3 moAb PR3G-2 as a gift. This manuscript has been released as a pre-print at bioRxiv (doi: 10.1101/559963).

## SUPPLEMENTARY MATERIAL

**Tables S1–S2 and Figure S1**

---

**Table S1.** Alpha carbon root mean square deviations (Å) among different PR3 variants.

|  | Xray-PR3-Ile<sup>103</sup> | Comp-PR3-Ile<sup>103</sup> | Comp-PR3-Val<sup>103</sup> | Com-iHm5-Val<sup>103</sup> |
| --- | --- | --- | --- | --- |
| Com-PR3-Ile<sup>103</sup> | 1.67 | 0 | 0.62 | 1.63 |
| Com-PR3-Val<sup>103</sup> | 1.90 | 0.62 | 0 | 1.56 |
| Com-iPR3-Val<sup>103</sup> | 1.75 | 0.59 | 0.45 | 1.41 |
| Com-iHm5-Val<sup>103</sup> | 2.34 | 1.63 | 1.56 | 0 |

**Table S2.**
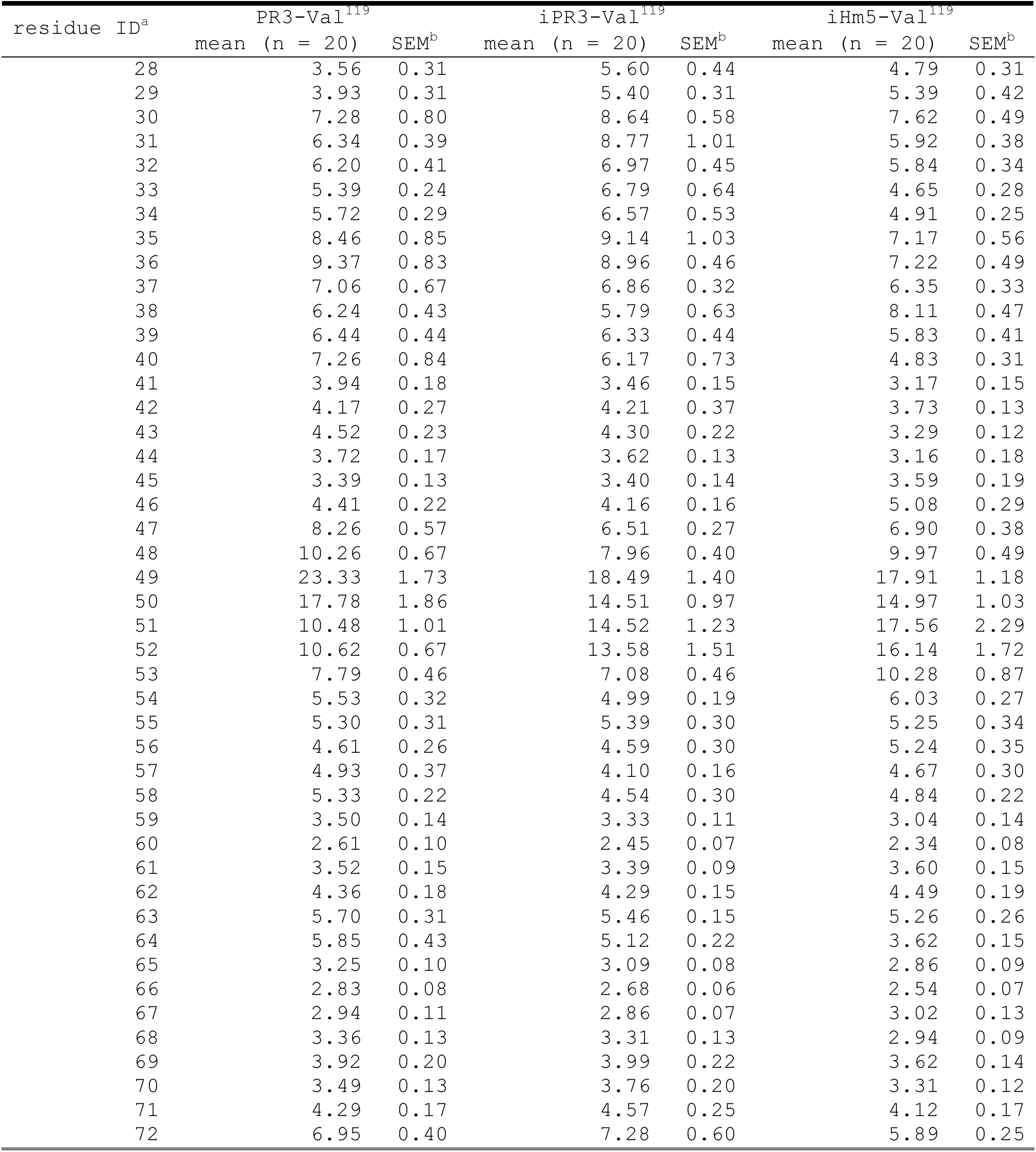
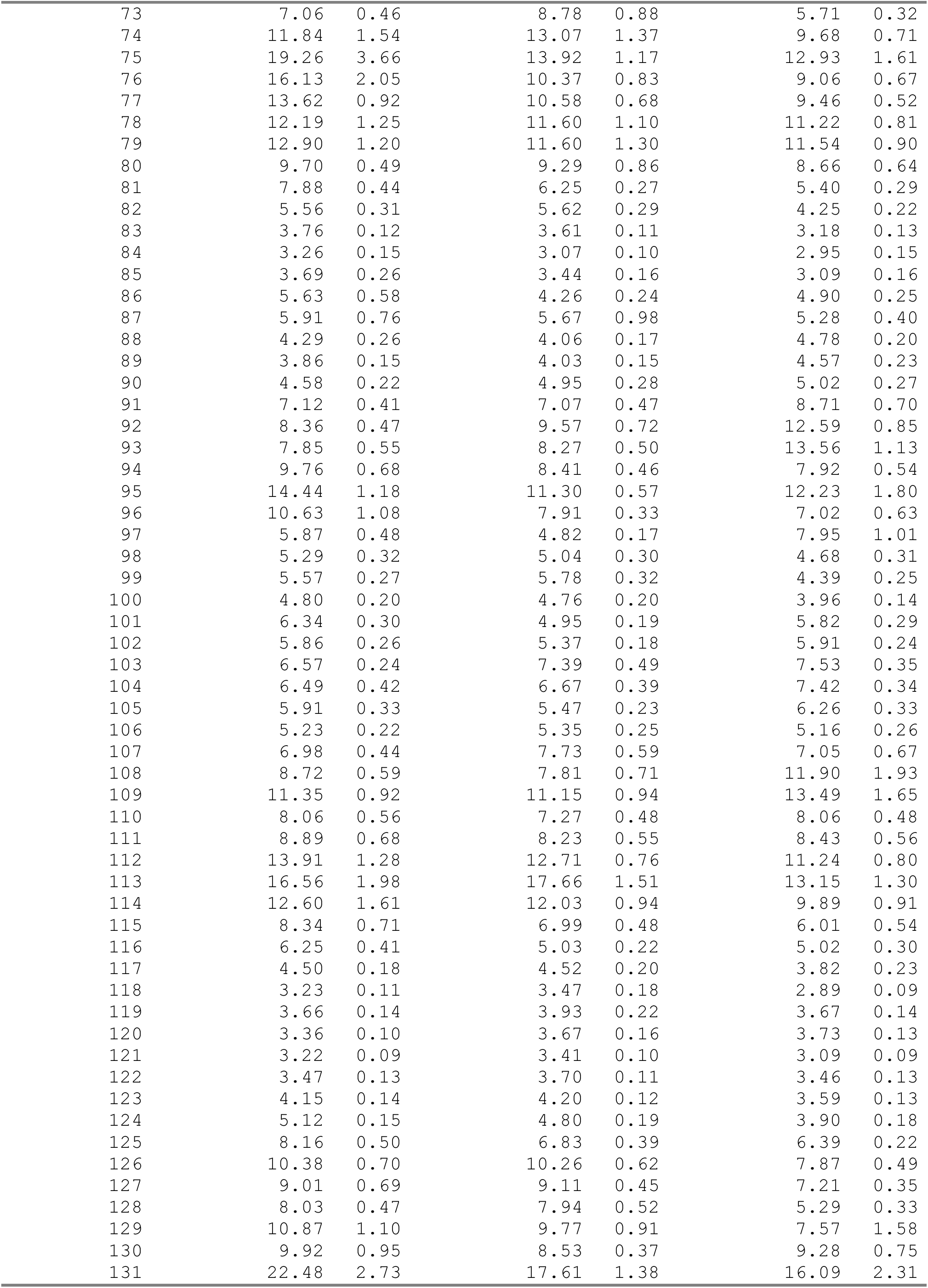
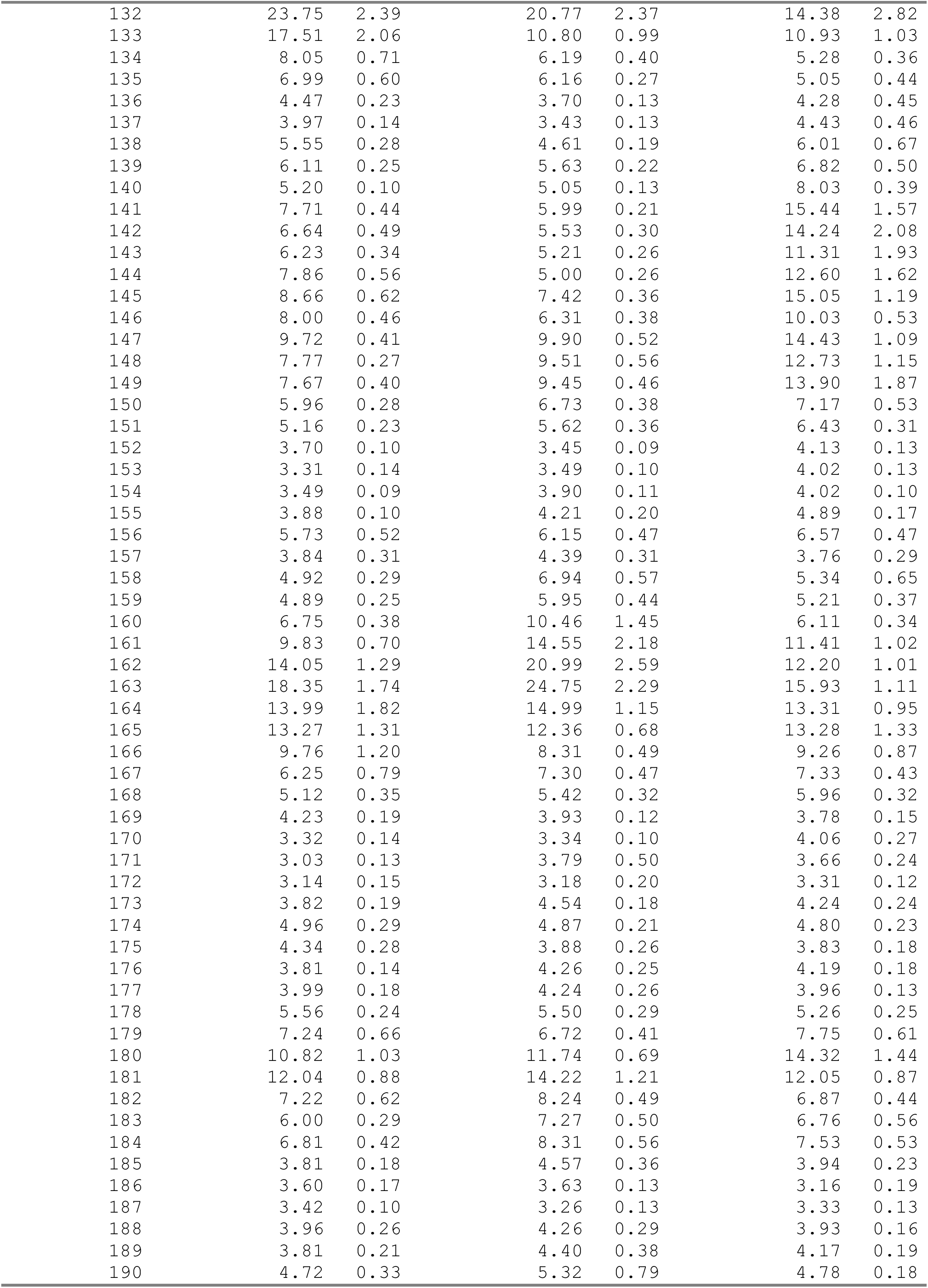
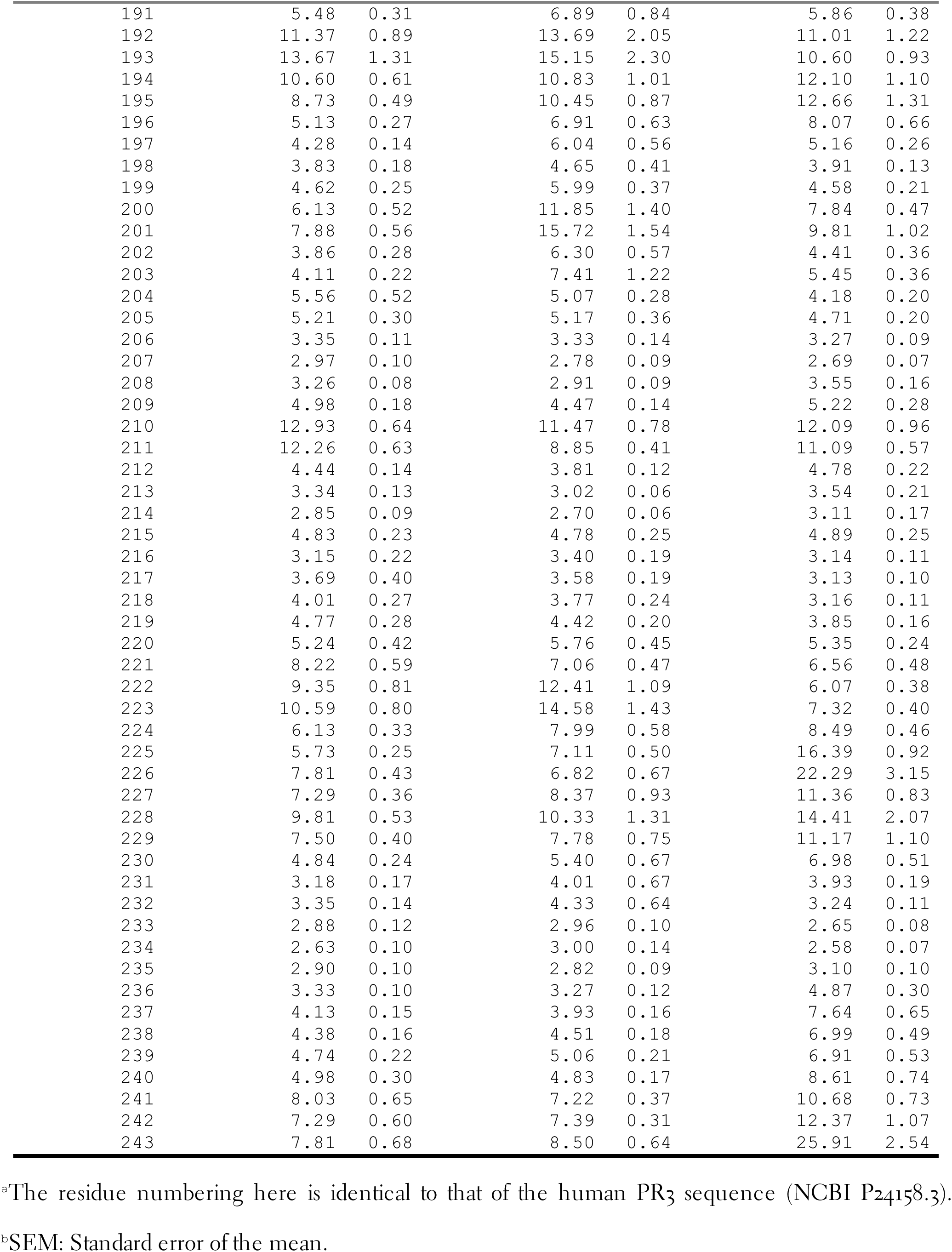
Alpha carbon B-factors of three PR3 variants.

**Figure S1.**
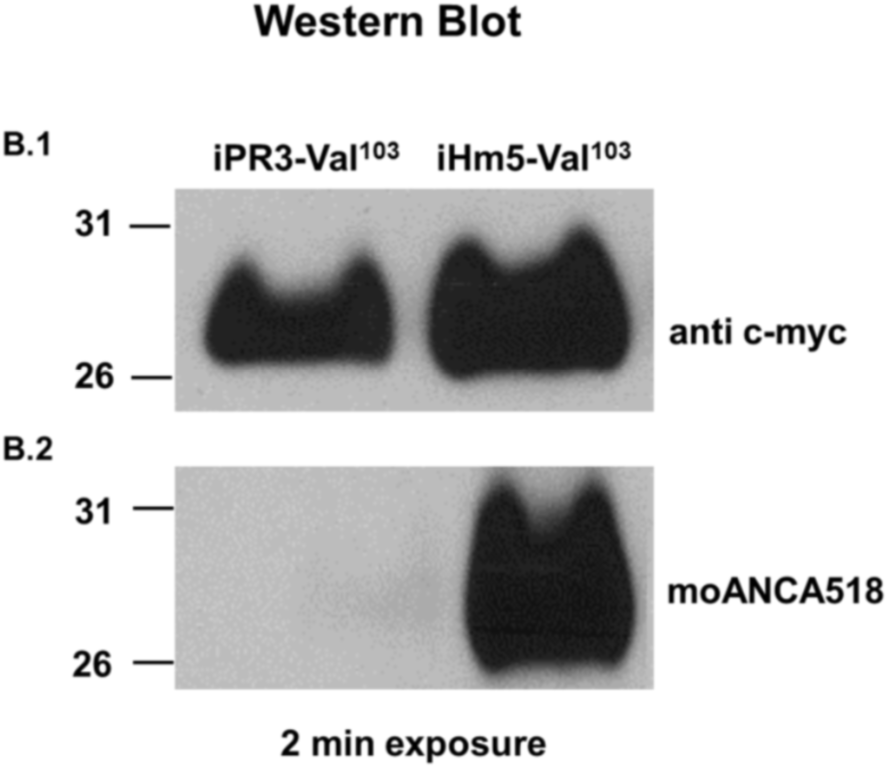
Western blots. B.1. Comparable binding of the murine anti-c-myc moAb (1.0 µg/mL) to the C-terminal cmyc-tag of the two antigens. B.2. Binding of moANCA518 (0.5 µg/mL) to iHm5-Val^103^ only.

## REFERENCES

1. Wegener F. Über generalisierte septische Gefäßerkrankungen. Verh Dtsch Pathol Ges (1936) 29:202–9.

2. Fahey JL, Leonard E, Churg J, Godman G. Wegener’s granulomatosis. Am J Med (1954) 17(2):168–79. doi: 10.1016/0002-9343(54)90255-7.

3. Fauci AS, Wolff SM. Wegener’s granulomatosis: studies in eighteen patients and a review of the literature. Medicine (Baltimore) (1973) 52(6):535–61. doi: 10.1097/00005792-197311000-00002.

4. Jenne DE, Tschopp J, Ludemann J, Utecht B, Gross WL. Wegener’s autoantigen decoded. Nature (1990) 346(6284):520. doi: 10.1038/346520a0.

5. Gross WL, Csernok E. Immunodiagnostic and pathophysiologic aspects of antineutrophil cytoplasmic antibodies in vasculitis. Curr Opin Rheumatol (1995) 7(1):11–9.

6. Gencik M, Meller S, Borgmann S, Fricke H. Proteinase 3 gene polymorphisms and Wegener’s granulomatosis. Kidney Int (2000) 58(6):2473–7. doi: 10.1046/j.1523-1755.2000.00430.x.

7. Specks U, Fass DN, Fautsch MP, Hummel AM, Viss MA. Recombinant human proteinase 3, the Wegener’s autoantigen, expressed in HMC-1 cells is enzymatically active and recognized by c-ANCA. FEBS Lett (1996) 390(3):265–70. doi: 10.1016/0014-5793(96)00669-2.

8. Sun J, Fass DN, Hudson JA, Viss MA, Wieslander J, Homburger HA, et al. Capture-ELISA based on recombinant PR3 is sensitive for PR3-ANCA testing and allows detection of PR3 and PR3-ANCA/PR3 immunecomplexes. J Immunol Methods (1998) 211(1-2):111–23. doi: 10.1016/S0022-1759(97)00203-2.

9. Sun J, Fass DN, Viss MA, Hummel AM, Tang H, Homburger HA, et al. A proportion of proteinase 3 (PR3)-specific anti-neutrophil cytoplasmic antibodies (ANCA) only react with PR3 after cleavage of its N-terminal activation dipeptide. Clin Exp Immunol (1998) 114(2):320–6. doi: 10.1046/j.1365-2249.1998.00730.x.

10. Specks U. What you should know about PR3-ANCA. Conformational requirements of proteinase 3 (PR3) for enzymatic activity and recognition by PR3-ANCA. Arthritis Res (2000) 2(4):263–7. doi: 10.1186/ar99.

11. Silva F, Hummel AM, Jenne DE, Specks U. Discrimination and variable impact of ANCA binding to different surface epitopes on proteinase 3, the Wegener’s autoantigen. J Autoimmun (2010) 35(4):299–308. doi: 10.1016/j.jaut.2010.06.021.

12. Capizzi SA, Viss MA, Hummel AM, Fass DN, Specks U. Effects of carboxy-terminal modifications of proteinase 3 (PR3) on the recognition by PR3-ANCA. Kidney Int (2003) 63(2):756–60. doi: 10.1046/j.1523-1755.2003.00765.x.

13. Lee AS, Finkielman JD, Peikert T, Hummel AM, Viss MA, Specks U. A novel capture-ELISA for detection of anti-neutrophil cytoplasmic antibodies (ANCA) based on c-myc peptide recognition in carboxy-terminally tagged recombinant neutrophil serine proteases. J Immunol Methods (2005) 307(1-2):62–72. doi: 10.1016/j.jim.2005.09.004.

14. Finkielman JD, Lee AS, Hummel AM, Viss MA, Jacob GL, Homburger HA, et al. ANCA are detectable in nearly all patients with active severe Wegener’s granulomatosis. Am J Med (2007) 120(7):643 e9–e14. doi: 10.1016/j.amjmed.2006.08.016.

15. Oommen E, Hummel A, Allmannsberger L, Cuthbertson D, Carette S, Pagnoux C, et al. IgA antibodies to myeloperoxidase in patients with eosinophilic granulomatosis with polyangiitis (Churg-Strauss). Clin Exp Rheumatol (2017) 35 Suppl 103(1):98–101.

16. Tan YC, Kongpachith S, Blum LK, Ju CH, Lahey LJ, Lu DR, et al. Barcode-enabled sequencing of plasmablast antibody repertoires in rheumatoid arthritis. Arthritis Rheumatol (2014) 66(10):2706– 15. doi: 10.1002/art.38754.

17. Tan YC, Blum LK, Kongpachith S, Ju CH, Cai X, Lindstrom TM, et al. High-throughput sequencing of natively paired antibody chains provides evidence for original antigenic sin shaping the antibody response to influenza vaccination. Clin Immunol (2014) 151(1):55–65. doi: 10.1016/j.clim.2013.12.008.

18. DeFalco J, Harbell M, Manning-Bog A, Baia G, Scholz A, Millare B, et al. Non-progressing cancer patients have persistent B cell responses expressing shared antibody paratopes that target public tumor antigens. Clin Immunol (2018) 187:37–45. doi: 10.1016/j.clim.2017.10.002.

19. Peikert T, Finkielman JD, Hummel AM, McKenney ME, Gregorini G, Trimarchi M, et al. Functional characterization of antineutrophil cytoplasmic antibodies in patients with cocaine-induced midline destructive lesions. Arthritis Rheum (2008) 58(5):1546–51. doi: 10.1002/art.23469.

20. Wiesner O, Russell KA, Lee AS, Jenne DE, Trimarchi M, Gregorini G, et al. Antineutrophil cytoplasmic antibodies reacting with human neutrophil elastase as a diagnostic marker for cocaine-induced midline destructive lesions but not autoimmune vasculitis. Arthritis Rheum (2004) 50(9):2954–65. doi: 10.1002/art.20479.

21. Wiesner O, Litwiller RD, Hummel AM, Viss MA, McDonald CJ, Jenne DE, et al. Differences between human proteinase 3 and neutrophil elastase and their murine homologues are relevant for murine model experiments. FEBS Lett (2005) 579(24):5305–12. doi: 10.1016/j.febslet.2005.08.056.

22. Van Der Geld YM, Limburg PC, Kallenberg CG. Characterization of monoclonal antibodies to proteinase 3 (PR3) as candidate tools for epitope mapping of human anti-PR3 autoantibodies. Clin Exp Immunol (1999) 118(3):487–96. doi: 10.1046/j.1365-2249.1999.01079.x.

23. Csernok E, Ludemann J, Gross WL, Bainton DF. Ultrastructural localization of proteinase 3, the target antigen of anti-cytoplasmic antibodies circulating in Wegener’s granulomatosis. Am J Pathol (1990) 137(5):1113–20.

24. Fujinaga M, Chernaia MM, Halenbeck R, Koths K, James MN. The crystal structure of PR3, a neutrophil serine proteinase antigen of Wegener’s granulomatosis antibodies. J Mol Biol (1996) 261(2):267–78. doi: 10.1006/jmbi.1996.0458.

25. Pang Y-P. FF12MC: a revised AMBER forcefield and new protein simulation protocol. Proteins (2016) 84(10):1490–516. doi: 10.1002/prot.25094.

26. Jorgensen WL, Chandreskhar J, Madura JD, Impey RW, Klein ML. Comparison of simple potential functions for simulating liquid water. J Chem Phys (1983) 79(2):926–35. doi: 10.1063/1.445869.

27. Pang Y-P. Use of 1–4 interaction scaling factors to control the conformational equilibrium between α-helix and β-strand. Biochem Biophys Res Commun (2015) 457(2):183–6. doi: 10.1016/j.bbrc.2014.12.084.

28. Berendsen HJC, Postma JPM, van Gunsteren WF, Di Nola A, Haak JR. Molecular dynamics with coupling to an external bath. J Chem Phys (1984) 81(8):3684–90. doi: 10.1063/1.448118.

29. Darden TA, York DM, Pedersen LG. Particle mesh Ewald: An N log(N) method for Ewald sums in large systems. J Chem Phys (1993) 98(12):10089–92. doi: 10.1063/1.464397.

30. Joung IS, Cheatham TE. Determination of alkali and halide monovalent ion parameters for use in explicitly solvated biomolecular simulations. J Phys Chem B (2008) 112(30):9020–41. doi: 10.1021/jp8001614.

31. Pang Y-P. Low-mass molecular dynamics simulation for configurational sampling enhancement: More evidence and theoretical explanation. Biochem Biophys Rep (2015) 4:126–33. doi: 10.1016/j.bbrep.2015.08.023.

32. Pang Y-P. At least 10% shorter C–H bonds in cryogenic protein crystal structures than in current AMBER forcefields. Biochem Biophys Res Commun (2015) 458(2):352–5. doi: 10.1016/j.bbrc.2015.01.115.

33. Andraos J. On the propagation of statistical errors for a function of several variables. J Chem Educ (1996) 73(2):150–4. doi: 10.1021/ed073p150.

34. Shao J, Tanner SW, Thompson N, Cheatham III TE. Clustering molecular dynamics trajectories: 1. Characterizing the performance of different clustering algorithms. J Chem Theory Comput (2007) 3(6):2312–34. doi: 10.1021/ct700119m.

35. Pang Y-P. How fast fast-folding proteins fold in silico. Biochem Biophys Res Commun (2017) 492(1):135–9. doi: 10.1016/j.bbrc.2017.08.010.

36. Van Regenmortel MH. Reductionism and the search for structure-function relationships in antibody molecules. J Mol Recognit (2002) 15(5):240–7. doi: 10.1002/jmr.584.

37. McCammon JA, Gelin BR, Karplus M. Dynamics of folded proteins. Nature (1977) 267(5612):585– 90. doi: 10.1038/267585a0.

38. Westhof E, Altschuh D, Moras D, Bloomer AC, Mondragon A, Klug A, et al. Correlation between segmental mobility and the location of antigenic determinants in proteins. Nature (1984) 311(5982):123–6. doi: 10.1038/311123a0.

39. Tainer JA, Getzoff ED, Alexander H, Houghten RA, Olson AJ, Lerner RA, et al. The reactivity of anti-peptide antibodies is a function of the atomic mobility of sites in a protein. Nature (1984) 312(5990):127–34. doi: 10.1038/312127a0.

40. Debye P. Interference of x rays and heat movement. Ann Phys (1913) 43:49–95.

41. Waller I. On the effect of thermal motion on the interference of X-rays. Z Phys (1923) 17:398–408.

42. Willis BTM, Pryor AW. Thermal vibrations in crystallography. London: Cambridge University Press (1975). 296 p.

43. Yu HA, Karplus M, Hendrickson WA. Restraints in temperature-factor refinement for macromolecules: An evaluation by molecular dynamics. Acta Crystallogr, Sect B: Struct Sci (1985) 41(Jun):191–201. doi: 10.1107/S0108768185001926.

44. Kidera A, Go N. Normal mode refinement: Crystallographic refinement of protein dynamic structure. 1. Theory and test by simulated diffraction data. J Mol Biol (1992) 225(2):457–75. doi: 10.1016/0022-2836(92)90932-A.

45. McRee DE. Practical protein crystallography. San Diego: Academy Press (1993). 386 p.

46. Trueblood KN, Burgi HB, Burzlaff H, Dunitz JD, Gramaccioli CM, Schulz HH, et al. Atomic displacement parameter nomenclature: Report of a subcommittee on atomic displacement parameter nomenclature. Acta Crystallogr, Sect A (1996) 52:770–81. doi: 10.1107/S0108767396005697.

47. Tronrud DE. Knowledge-based B-factor restraints for the refinement of proteins. J Appl Crystallogr (1996) 29:100–4. doi: 10.1107/S002188989501421x.

48. Garcia AE, Krumhansl JA, Frauenfelder H. Variations on a theme by Debye and Waller: From simple crystals to proteins. Proteins (1997) 29(2):153–60. doi: 10.1002/(SICI)1097-0134(199710)29:2<153::AID-PROT3>3.0.CO;2-E.

49. Artymiuk PJ, Blake CC, Grace DE, Oatley SJ, Phillips DC, Sternberg MJ. Crystallographic studies of the dynamic properties of lysozyme. Nature (1979) 280(5723):563–8. doi: 10.1038/280563a0.

50. Morin S. A practical guide to protein dynamics from ^15^N spin relaxation in solution. Prog Nucl Magn Reson Spectrosc (2011) 59(3):245–62. doi: 10.1016/j.pnmrs.2010.12.003.

51. Pang Y-P. Use of multiple picosecond high-mass molecular dynamics simulations to predict crystallographic B-factors of folded globular proteins. Heliyon (2016) 2(9):e00161. doi: 10.1016/j.heliyon.2016.e00161.

52. Hinkofer LC, Seidel SA, Korkmaz B, Silva F, Hummel AM, Braun D, et al. A monoclonal antibody (MCPR3-7) interfering with the activity of proteinase 3 by an allosteric mechanism. J Biol Chem (2013) 288(37):26635–48. doi: 10.1074/jbc.M113.495770.

53. Thompson GE, Casal Moura M, Nelson DA, Hummel A, Jenne DE, Fervenza FC, et al. Characterization of preferential recognition of a chimeric recombinant proteinase 3 variant by anti-neutrophil cytoplasmic antibodies. Arthritis Rheumatol; Chicago(2018).

54. Travers TS, Harlow L, Rosas IO, Gochuico BR, Mikuls TR, Bhattacharya SK, et al. Extensive Citrullination Promotes Immunogenicity of HSP90 through Protein Unfolding and Exposure of Cryptic Epitopes. J Immunol (2016) 197(5):1926–36. doi: 10.4049/jimmunol.1600162.

